# Acute resistance to BET inhibitors remodels compensatory transcriptional programs via p300 co-activation

**DOI:** 10.1101/2022.09.14.507850

**Authors:** Viral Shah, George Giotopoulos, Hikari Osaki, Markus Meyerhöfer, Eshwar Meduri, Benedict Schubert, Haiyang Yun, Sarah J Horton, Shuchi Agrawal-Singh, Patricia S Haehnel, Faisal Basheer, Dave Lugo, Michael WM Kühn, Borhane Guezguez, Matthias Theobald, Thomas Kindler, Paolo Gallipoli, Rab K Prinjha, Brian JP Huntly, Daniel Sasca

**Author notes:** Shared first authors (Viral Shah, George Giotopoulos, Hikari Osaki). Co-senior and Corresponding authors.

## Abstract

Initial clinical trials with drugs targeting epigenetic modulators - such as bromodomain and extraterminal (BET) inhibitors - demonstrate modest results in acute myeloid leukemia (AML). The main reason for this involves an increased transcriptional plasticity within AML, which allows cells to escape the therapeutic pressure. To study mechanisms of resistance, we investigated immediate epigenetic and transcriptional responses following BET inhibition, and could demonstrate that BET inhibitor-mediated release of BRD4 from chromatin is accompanied by an acute compensatory feedback loop that attenuates inhibition, or even increases expression, of specific transcriptional modules. This adaptation is most marked at key AML maintenance genes and is mediated by p300, suggesting a rational therapeutic opportunity by combining BET- and p300- inhibition. p300 activity is required during all steps of adaptation. However, the transcriptional programs that p300 regulates to induce resistance to BETi differ between AML subtypes. Remarkably, in some AMLs, p300 regulates a series of transitional transcriptional patterns that allow homeostatic adjustments during earlier stages of resistance to BET-inhibitors. In consequence, p300 remains crucial throughout all stages of resistance in sensitive AML-subtypes, although its importance declines following the development of chronic resistance to BET inhibitors in some other AMLs. Altogether, our study elucidates the mechanisms that underlie an “acute” state of resistance to BET inhibition, achieved through p300 activity, and how these mechanisms remodel to become “chronic”. Importantly, however, our data also suggest that a sequential treatment with BET- and p300 inhibition may prevent resistance development, thereby improving outcomes.

**Key points:**

- A mechanistic feedback to p300 enables acute tolerance to BET inhibition.
- p300 regulates transcriptional networks that lead to chronic resistance to BET inhibition.
- Sequential BET-, followed by p300-inhibition, is synthetically lethal in AML, and is optimally deployed during earlier stages of resistance to BET inhibitors.

## Introduction

Acute myeloid leukemias (AML) are aggressive cancers caused by an increased capacity of hematopoietic progenitors to proliferate, block cellular maturation and adapt to treatment, characteristics often dependent upon an increased cellular “plasticity”^1^. Dysregulated epigenetic and transcriptional processes are central determinants of this AML plasticity^2–4^. As a result, modulators of aberrant transcriptional activation are, theoretically, attractive therapeutic targets, with bromodomain and extraterminal protein (BET) inhibitors serving as paradigmatic novel epigenetic treatments in AML^5–7^. BET inhibitors predominantly target BRD4, an epigenetic reader that recognizes acetylated histones H3/H4 and transcription factors (TF), mediates transcriptional elongation and enhancer-directed transcription and further serves as a scaffold for the recruitment of other co-factor proteins^8–11^. Although BRD4 binds chromatin genome-wide, its common association at cancer-maintenance genes (e.g. *MYC* or *RUNX1*) decorated with high-level acetylation leads to the selective dependence of cancers on BRD4 and to its therapeutic window of inhibition^12^.

However, a major issue with BET inhibitors became apparent after their introduction into clinical trials. This consisted of modest response rates in monotherapy due to either primary or acquired resistance^13–18^. As has been demonstrated for resistance to other novel epigenetic drugs, BET inhibitor resistance appears predominantly non-genetic, suggesting that epigenetic adaptation underlies this resistance^13,17–21^. This is further evidenced by the rapid restoration of *MYC* transcription in BET inhibitor-resistant cells despite continued inhibition^18^, or by recent reports about treatment strategies to target BET inhibitor-resistant cells, using either epigenetic inhibitors to LSD1 or CDK7^13,22^.

AML exemplifies a group of diseases that usually respond to initial treatment but later relapse, often fatally. The development of resistance requires both the initial survival and perseverance of a population of cells under the selective pressure of therapy^23,24^. However, the mechanisms that underpin these acute processes are poorly understood. Furthermore, it remains unexplained as to whether mechanisms underpinning initial adaptation and later full-blown resistance are uniform or further evolve. Obtaining such knowledge will allow better design of upfront treatment and possible maintenance strategies.

Here, we hypothesize that transcriptional plasticity in AML mediates early pathways of adaptation to and escape from BET inhibition. Moreover, we predict that this plasticity also provides new epigenetic vulnerabilities that, when properly inhibited, may extinguish later resistance. To test this, we experimentally deconvoluted the acute alterations of BRD4-associated transcription in two sensitive but mechanistically disparate subtypes of AML, following BET inhibition in initially sensitive AML. This allowed us to identify an acute reliance on p300 activity to rescue critical leukemia maintenance programs, which became functionally evident in a broader range of AMLs upon testing. We further charted longitudinal alterations in expression with the transition from acute to chronic resistance that demonstrate a dynamic but continued requirement for p300-mediated transcription, along with rewired transcription to maintain critical leukemic programs.

## Methods

### Establishment of BETi-resistant cells

To generate longitudinal BETi resistance models, OCI-AML3 (harbouring *NPMlc, DNMT3A, SPI1, NSD3* and *KRAS* mutations) (RRID:CVCL_1844) and SKNO1 (harbouring an *RUNX1-RUNXT1* gene rearrangement and a *C-KIT* activating mutation) (RRID: CVCL_2196) cells were plated in 24-well plates and continuously exposed to increasing concentrations of BETi (for SKNO1: IC20-40 nM BETi, IC40-50 nM BETi, IC50-100 nM BETi, IC65-175 nM BETi, IC90-250 nM BETi; for OCI-AML3: IC40-100 nM BETi, IC50-150 nM BETi, IC65-200 nM BETi and IC90-300 nM BETi) for at least 4 weeks per each concentration step. At least 3 clones were then expanded and grown for 2 more weeks at each resistance stage, then partly aliquoted and frozen down for later use and partly re-plated into 24-well plates under continuous treatment with the next-higher concentration. DMSO-treated clones were maintained identically.

### Data availability

All RNASeq, ChIPSeq, ATACSeq and nucRNASeq data have been deposited in the GEO database under the accession number GSE167163 and will be released upon publication.

## Results

### RUNX1-RUNX1T1 leukemia as a model of BET dependency

To probe early responses to BET inhibition in relevant leukemic models, we defined a set of 6 cell lines representative of the genetic heterogeneity of AML, but with the common characteristic of sensitivity to BET inhibition (Supplementary Table S1). We treated these cells with different concentrations of iBET-151 (hereafter BETi), a tool compound we have previously demonstrated to be pharmacologically matched to Molibresib^5^ (Figure 1A). We and others have previously presented extensive functional effects of BET inhibition in MLL-rearranged or NPM1-mutated AML^5,7,12^. In addition, complementary to previous data, our experiments demonstrated that cell lines that carry the t(8;21) translocation and rearrange *RUNX1-RUNX1T1* (KASUMI-1 and SKNO-1) conferred the highest sensitivity to BETi^25^. BETi rapidly induced cell death in both *RUNX1-RUNX1T1-positive* cell lines (Figure 1B). Furthermore, BETi significantly reduced colony numbers (Figure 1C and S1A), results that were confirmed in 2 separate RUNX1-RUNX1T1-driven patient samples (Figure 1D).

**Figure 1.**
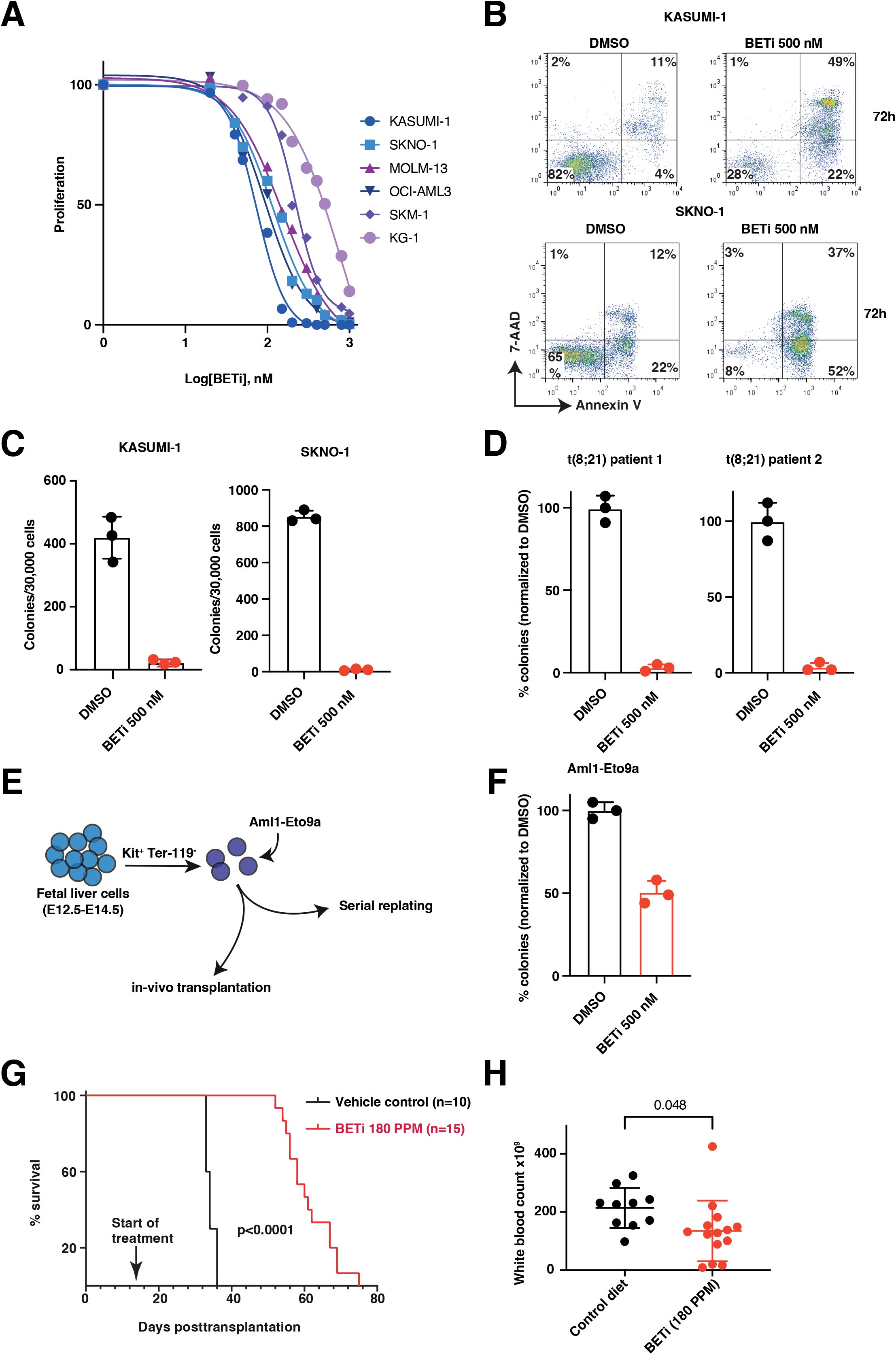
RUNX1;RUNX1T1 leukemia as a model of BET dependency. A. Analysis of cell proliferation of the indicated BETi-sensitive AML cell lines after 72h of BETi treatment. Shown are mean percentages normalized to DMSO-treated controls and standard deviations (SD) from 3 biological replicates. B. Representative scatter plots showing Annexin V/7-AAD at 72h after treatment of KASUMI-1 and SKNO1 cells with either DMSO or BETi. C. Assessment of colony formation of KASUMI-1 and SKNO1 cells after treatment with either DMSO or BETi for 7 days in methylcellulose. Shown are mean colony numbers and SD from 3 biological replicates. D. Assessment of colony formation of 2 different t(8;21) AML primary patient samples after treatment with either DMSO or BETi for 7 days in methylcellulose. Shown are mean percentages normalized to DMSO and SD from 3 biological replicates. E. Schematics of procedures to produce a model of murine *Am/1-Eto9a-driven* AML and analyze its sensitivity to BETi. F. Analysis of colony formation of *Am/1-Eto9a*-transformed murine AML cells after treatment with either DMSO or BETi for 7 days in methylcellulose. Shown are mean percentages normalized to DMSO and SD from 3 biological replicates. G. Kaplan-Meier survival curves of secondary recipient animals transplanted with equal numbers of *Am/1-Eto9a-driven* murine AML cells and orally treated with chow food containing the indicated compounds. PPM = parts per million in chow food, equivalent to approximately 30 mg/kg/d ingested BETi. Differences were compared using the log-rank test. H. Analysis of white blood cell counts (WBC) at terminal endpoints between control and BETi diet mice from Figure 1G.

To confirm the functional role of BETi in t(8;21) AML *in-vivo,* we established a previously described *Aml1-Eto9a-driven* murine AML^26^. cKIT-high/Ter-119-negative murine fetal liver cells were transduced with retroviral particles containing *Aml1-Eto9a* and either transplanted into mice or serially plated in methylcellulose (Figure 1E). Transformation occurred in both settings (Figures S1B-D). *In-vitro* transformed cells were treated with BETi or DMSO, resulting in a significant reduction of colony numbers for the BETi-treated arm (Figure 1F). As expected, *in-vivo* treatment with BETi significantly increased disease latency and improved overall survival (Figure 1G-H). Altogether, these data demonstrate that BET proteins are critical therapeutic targets in *RUNX1-RUNX1T1* AML.

### Exclusion of BRD4 from chromatin via BET inhibition induces redistribution of p300 to critical AML maintenance genes

Leukemic maintenance in t(8;21) AML relies on a refined balance between RUNX1-RUNX1T1 and the wild-type RUNX1 protein^27–31^. Cooperative binding of PU.1, FLI1, ERG, and LMO2 ^29,32^ further influences this balance, while p300/CREBBP mediates local activation through acetylation of lysine residues of histone tails and TFs^33–35^. Acetylation recruits BRD4 that, together with the PTEF-b complex, facilitates transcriptional elongation at paused promoters^36,37^. As the mechanisms underlying t(8:21) AML transformation and maintenance are described in great detail, this model was deemed optimal to deconvolute early chromatin processes upon BETi. We first analyzed KASUMI-1 cells at 24h post-treatment. This was long enough to allow release of BRD4 from chromatin, but prior to the onset of cell-cycle arrest/death (Figure 2B and S2A-B). We assessed binding of the above disease drivers and regulators (RUNX1, RUNX1T1 (ETO), PU.1, FLI1, ERG, LMO2, p300, CDK9) and H3K27ac via chromatin immunoprecipitation followed by massive parallel sequencing (ChIP-Seq). To link these with dynamics of chromatin accessibility and transcription, we also measured differential nuclear RNASeq (nucRNASeq) and ATACSeq (overview in Figure 2A).

**Figure 2.**
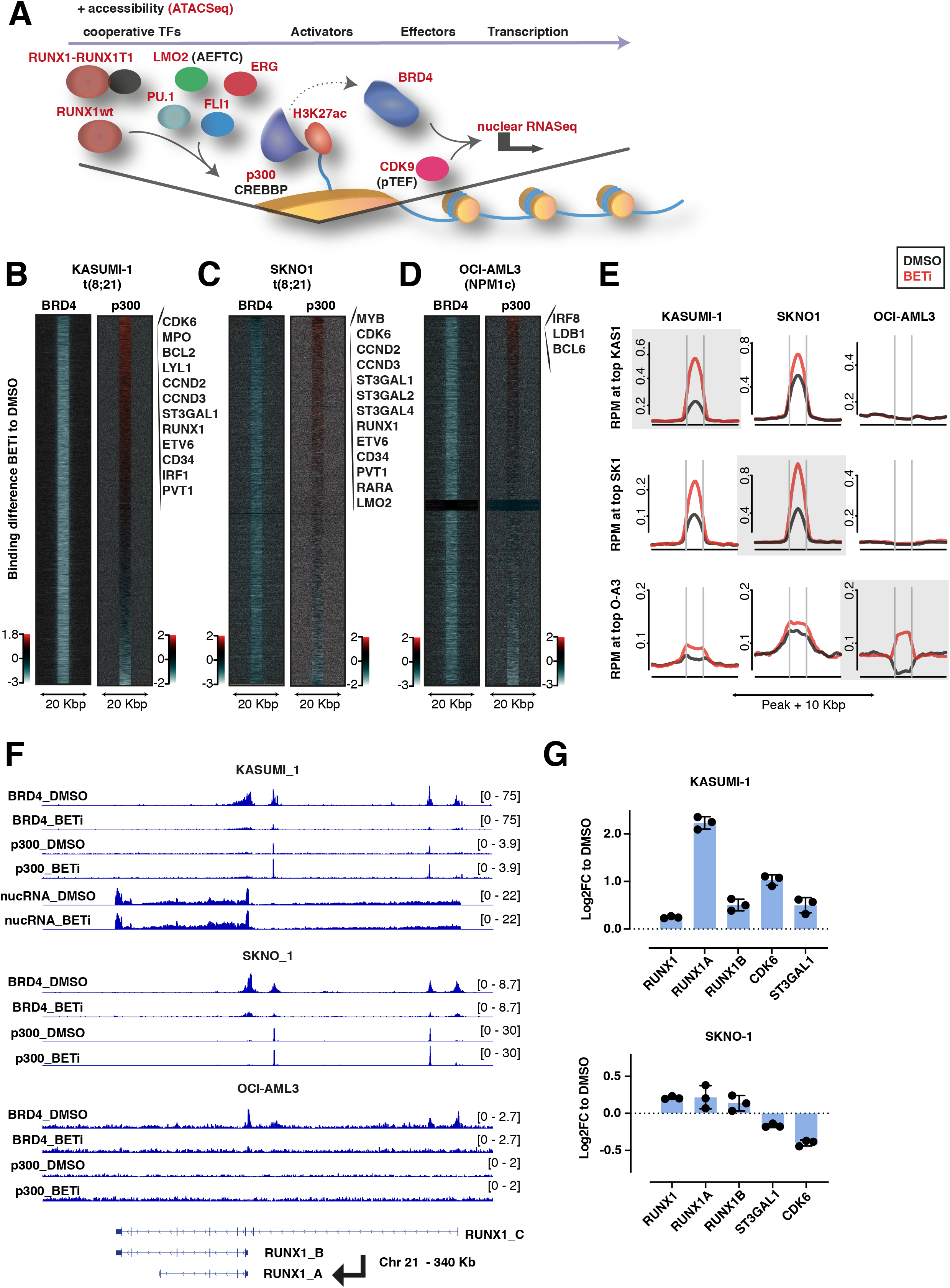
Removal of BRD4 from chromatin via BET inhibition induces redistribution of p300 to critical AML maintenance genes. A. Schematic depicting the experimental strategy to determine dynamics at BRD4-bound chromatin after 24h of DMSO or BETi treatment in KASUMI-1 cells. In t(8:21), RUNX1-RUNX1T1 and RUNX1 compete for binding to chromatin. In doing so, they recruit further cooperative TFs such as LMO2, PU.1, FLI1 and ERG. To “activate” transcription, they rely on the transcriptional co-activators p300 and CREBBP, which acetylate H3K27ac and recruit BRD4. In conjunction with pTEF (CDK9), BRD4 leads the transcriptional elongation. ChIPSeq was performed at 24h of treatment with either DMSO or BETi for all factors marked with red, and also for ATACSeq and nucRNASeq. For the initial pattern identification, single replicates per condition were acquired for ChIPSeq profiles and 2 biological replicates per condition for nucRNASeq and ATACSeq. B.-D. Tornado plots of BRD4 and p300 binding differences between BETi and DMSO conditions in KASUMI-1, SKNO1 and OCI-AML3 cells (24h treatment). Positive enrichment (in red) shows stronger binding upon BETi. Negative enrichment, colored in white, shows loss of binding upon BETi. Shown are representative matched replicates. With the exception of BRD4 ChIPSeq in KASUMI-1, which was performed once per condition, all experiments were performed with 2 biological replicates per condition. E. Average binding curve profiles for p300 at the top 5% rescued sites in the indicated cell lines. Upper panels show binding at the 5% best-scoring sites that were found in KASUMI1, middle panels show binding profiles at the 5% best-scoring sites that were found in SKNO1, lower panels apply the same method for 5% best-scoring sites that were found in OCI-AML3. F. Examples of BRD4 and p300 binding profiles in DMSO and BETi-treated KASUMI-1 (here also nucRNASeq dynamics), SKNO1 and OCI-AML3 cells, to demonstrate the BETi-triggered increase of p300 binding (and rescue of mRNA production) at the indicated genomic loci. G. Analysis of qPCR expression of the indicated transcripts/genes in KASUMI-1 and SKNO1 cells after treatment with either DMSO or BETi (24h). Shown are log2 Fold Changes normalized to DMSO-treated controls and SD from 3 biological replicates.

As expected, BRD4 was significantly excluded from chromatin by BETi (Figure 2B) and disruption was strongest at/near critical AML maintenance genes that demonstrated strong basal BRD4 binding, as exemplified by *CDK6, BCL2, CCND2-3, ST3GAL1, RUNX1, ETV6, CD34, PVT1* and other loci (Figure 2B). Strikingly, it was obvious that p300 binding to many of these critical AML maintenance genes significantly increased after BET inhibition (Figure 2B). However, although a global decrease in binding was observed for ERG (that was also accompanied by a decreased *ERG* expression) no significant differential binding was evident for RUNX1, FLI1, RUNX1-RUNX1T1, LMO2, SPI1, CDK9 and H3K27ac (Figure S2C). In addition, no significant global differences were noted for chromatin accessibility (ATACSeq) and mRNA abundance (nuclear RNASeq) (Figure S2C).

We hypothesized that the increased p300 binding may have occurred as a feedback mechanism to rescue critical AML maintenance genes that were normally dependent upon the transcriptional effector BRD4. To observe whether the locus-specific accumulation of p300 after BET inhibition was reproducible in other AML cell lines, we performed ChIP-Seq for BRD4 and p300 using the same conditions (at 24h following either DMSO or BETi treatment) in SKNO1 (another t(8;21) cell line) and in a completely independent but BETi-sensitive AML cell line, OCI-AML3 (driven by mutated *NPM1).* In SKNO1 cells, a similar marked increase in binding of p300 became evident at sites corresponding to those in KASUMI-1 (as exemplified by *CDK6, CCND2-3, ST3GAL1-2, 4, RUNX1, ETV6, CD34* and *PVT1*) (Figure 2C). In slight contrast, in OCI-AML3 cells, although we observed an increase of p300 binding at distinct loci, these were fewer and exemplified by a different set of genes including *IRF8, LDB1* and *BCL6* (Figure 2D).

We also compared signal intensities across all 3 models at the top 5% sites of increased p300 binding in each cell line. Upon BETi, p300 signal intensities at these putative “rescue” sites in KASUMI-1 were likewise significantly higher in SKNO1 and vice-versa. However, OCI-AML3 cells had a unique pattern, with a generally low p300 binding to chromatin altogether, and with no overlap with the top 5% from KASUMI-1 or SKNO1. Still, there was an obvious new acquisition of p300 at a number of loci in the enforced absence of BRD4 (Figure 2E bottom-right). Representative examples for the p300-mediated putative rescue can be observed in Figures 2F and S2E-F. Upon BETi, BRD4 is lost from chromatin and p300 is gained at the same binding sites in KASUMI-1 and SKNO-1 (Figure 2F and S2E) or OCI-AML3 (Figure S2F showing putative rescue at an IRF8 enhancer) cell lines.

Comparing differential binding of the regulators shown in Fig 2A at the top 5% of KASUMI-1 putative rescue sites, we observed significant enrichment for binding of RUNX1, RUNX1-RUNX1T1 and FLI1, however only marginally higher signal intensity for H3K27ac (Figure S2D). All other parameters showed either no change or a decrease in binding upon BETi (data not shown).

Notably, the nucRNASeq signal in KASUMI-1 surprisingly did not decreased (but instead slightly increased) at the major RUNX1 isoforms -A and -B (Figure 2F). As we have observed milder than expected decreases of mRNA abundance at more such putative rescue sites (data not shown due to redundancy with qPCR), we verified the real-time mRNA production in KASUMI-1 and SKNO1 cells via qPCR for RUNX1 and separately for the RUNX1 isoforms -A and -B, and also for the putative rescue genes ST3GAL1 and CDK6. After 24h with BETi, both RUNX1 isoforms and the “full” product were expressed higher than at baseline, with CDK6 and ST3GAL1 also higher in KASUMI-1 and only slightly reduced in SKNO1 cells (Figures 2G and S2E).

Altogether, these results suggest an unanticipated compensatory maintenance of critical genes following BETi, whereby redistribution of p300 to “rescue” loci blunts the downregulation of these critical leukemia effectors. While more obvious in t(8:21) AML, this acute compensation occurs across different AML subtypes and is subtype-specific.

### Sequential pharmacological BET-followed by p300-inhibition synergistically/optimally suppresses AML proliferation

These data suggest an immediate compensatory mechanism to maintain transcription of several critical genes via p300 upon BETi (Figure 3A). To determine if this acute response to BETi also provided a therapeutic vulnerability and therefore opportunity, we used a recently described lysine acetyltransferase inhibitor with high potency and selectivity for p300/CREBBP - A-485 (hereafter p300i)^38^. We treated KASUMI-1, SKNO1 and OCI-AML3 cells with the two inhibitors in 3 temporal sequences: mode i) BETi sequentially followed by p300i, mode ii) both inhibitors concomitantly and mode iii) p300i sequentially followed by BETi. Treatment was performed using BETi and p300i each at 4 dosages + DMSO controls to a total of 20 combinations. This strategy allowed for an unbiased analysis of treatment effects and the determination of synergistic activity, as quantified using the ZIP calculation algorithm^39^. We defined a ZIP threshold of +10 for synergism. Importantly, BETi+p300i treatment was invariably synergistic only in mode i) BETi-first (Figure 3B-D). KASUMI-1 and SKNO1, but not OCI-AML3, were synergistic in mode ii) (concomitant), while only SKNO1 cells remained synergistic in the p300i-first sequence (mode iii)) (Figure 3B-D).

**Figure 3.**
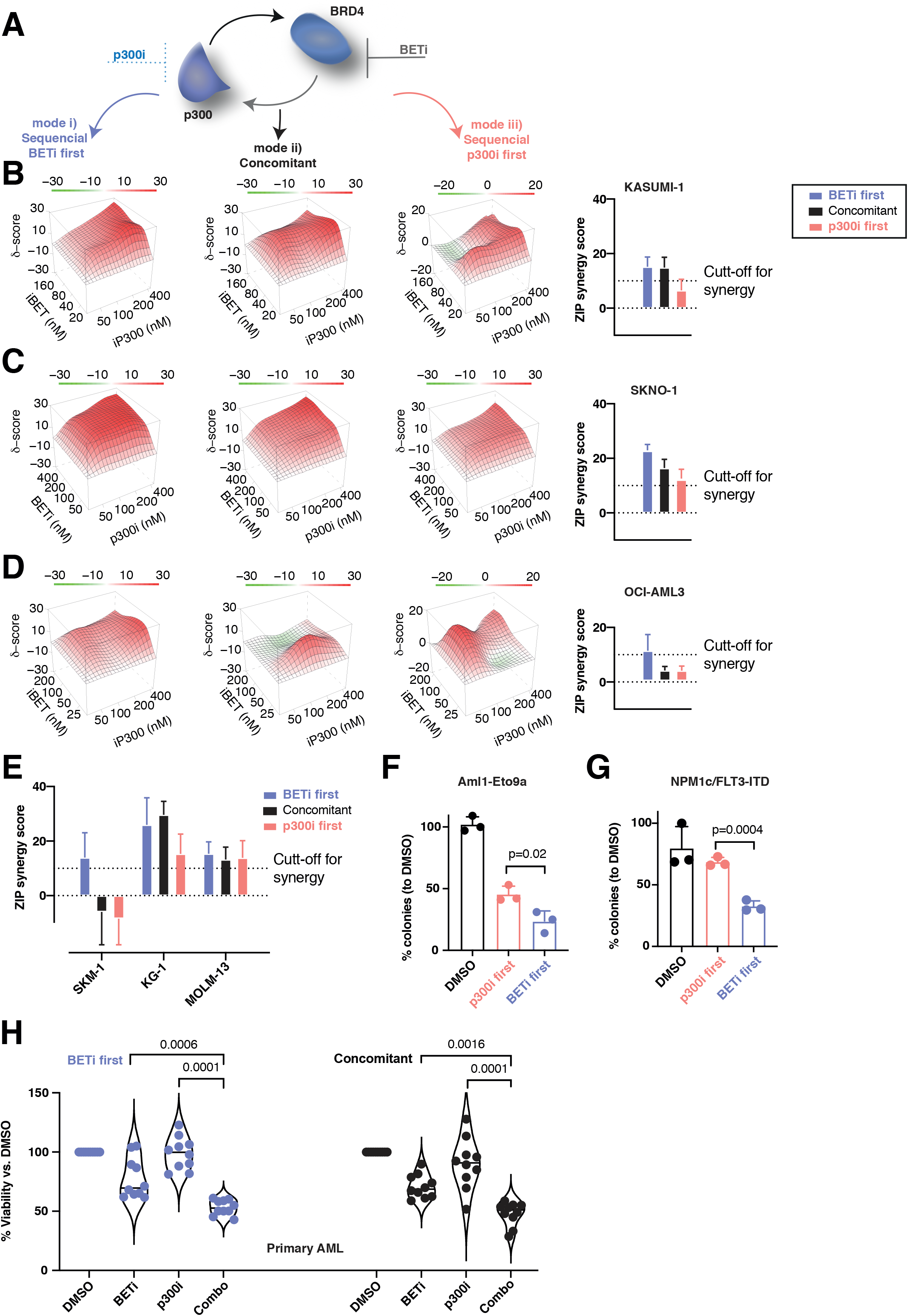
Sequential pharmacological BET-followed by p300-inhibition synergistically/optimally suppresses AML proliferation. A. Schematical approach of inhibition of the proposed feedback rescue loop to p300 after BETi. In total, 6 cells lines were treated in each 3 biological replicates with the two inhibitors at each 4 dosages and DMSO (20 combinations) in 3 temporal sequences: mode i) BETi sequentially followed by p300i, mode ii) both inhibitors concomitantly and mode iii) p300i sequentially followed by BETi. B.-D. Three-dimensional diffusion plots and plots of synergy scores (right panels) of combined treatment with BETi and p300i in the indicated orders and cell lines. Treatment efficiency was measured with CellTiterGlo^®^ at day 5 after begin of the first treatment. For the sequential treatment modes, the second compound was added 48 hours after treatment commencement. Shown are averages and (for bar plots) standard deviation from 3 biological/experimental replicates. E. Plots of synergy scores of combined treatment with BETi and p300i in the indicated orders and cell lines. Treatment efficiency was measured with CellTiterGlo^®^ at day 5 after begin of the first treatment. For the sequential treatment modes, the second compound was added 48 hours after treatment begin. Shown are averages and standard deviation from 3 biological/experimental replicates. F. Analysis of colony formation of Aml1-Eto9a-transformed murine AML cells after 2 rounds of plating and treatment with either DMSO in both plates, BETi followed by p300i or vice versa. Shown are mean percentages normalized to DMSO and SD from 3 biological replicates. In each plating, treatment was performed for 7 days in methylcellulose. G. Analysis of colony formation of NPM1c/FLT3-ITD murine AML cells after 2 rounds of plating and treatment with either DMSO in both plates, BETi followed by p300i or vice versa. Shown are mean percentages normalized to DMSO and SD from 3 biological replicates. In each plating, treatment was performed for 7 days in methylcellulose. H. Violin plots showing treatment efficacy of concomitant or sequential treatment with BETi and p300i in 5 primary AML patient samples. Shown are results from 2 biological replicates for each primary sample. For the sequential treatment mode with BETi first, p300i was added 48 hours after treatment commencement.

To assess the effects of combined BETi+p300i treatment across other AML genotypes, we extended the same analysis to more BETi sensitive cells - SKM1, MOLM-13 and KG-1. Again, synergistic inhibition of proliferation was present in all cell lines when utilizing mode i) the BETi first - p300i second treatment sequence (Figure 3E and S3A). Inhibition became less synergistic using concomitant treatment in SKM1, while only KG-1 and MOLM-13 demonstrated synergism when p300i was used ahead of BETi (Figure 3E and S3A). We also assessed for synergism in our models of *Am/1-Eto9a* and *Npm1*^flox-cA/+^;*Flt3*^ITD/+^;Mx1-Cre+ (*Npm1c/Flt3-ITD*)^4,40^ murine leukemic cells, analyzing colony replating capacity as a readout. Cells were replated for 2 weeks after sequential treatment with either BETi followed by p300i or vice-versa. While both sequences demonstrated reduced colony numbers when compared with DMSO, BETi followed by p300i treatment was significantly more effective than the same combination in the reverse sequence (Figure 3F-G). Finally, we confirmed our findings in a series of 5 patient samples *ex-vivo,* in which sequential BETi followed by p300i treatment demonstrated significant inhibition of proliferation to a comparable degree with the concomitant treatment mode (Figure 3H and S3B). In summary, these results demonstrate an increased therapeutic efficacy when BETi and p300i are combined. In keeping with our demonstration of a p300-mediated mechanism that perpetuates transcriptional programs necessary for the continued survival of BETi-treated cells, the combination was most efficient when cells were treated sequentially with BETi first.

### Sequential BETi p300i treatment counteracts expression of rescue programs in a cell type-specific manner

Given the overlap of some critical AML maintenance genes with p300-rescued genes during the acute adaptation phase to BETi, we hypothesized that BETi-dependent transcripts critical for AML maintenance are likewise subject to p300i-mediated inhibition. To demonstrate this, we performed RNASeq in our two t(8:21) cell lines (KASUMI-1 and SKNO1), and also in OCI-AML3 and MOLM-13 (a BETi-senstive t(9; 11) AML cell line that shares some transcriptional similarity with OCI-AML3) cells, after 24h of treatment with p300i or DMSO. In KASUMI-1 and SKNO1 cells, we observed a significant decrease of several p300-rescued genes, including *RUNX1, ST3GAL1* and *4, ETV6 or MYC* (Volcanos Figure 4A-B). Meanwhile, in OCI-AML3 and MOLM-13 cells, only a decrease of *MYC* expression was consistent, while p300i generally increased expression of standard interferon-stimulated genes (ISGs) (Volcanos Figure 4C-D).

**Figure 4.**
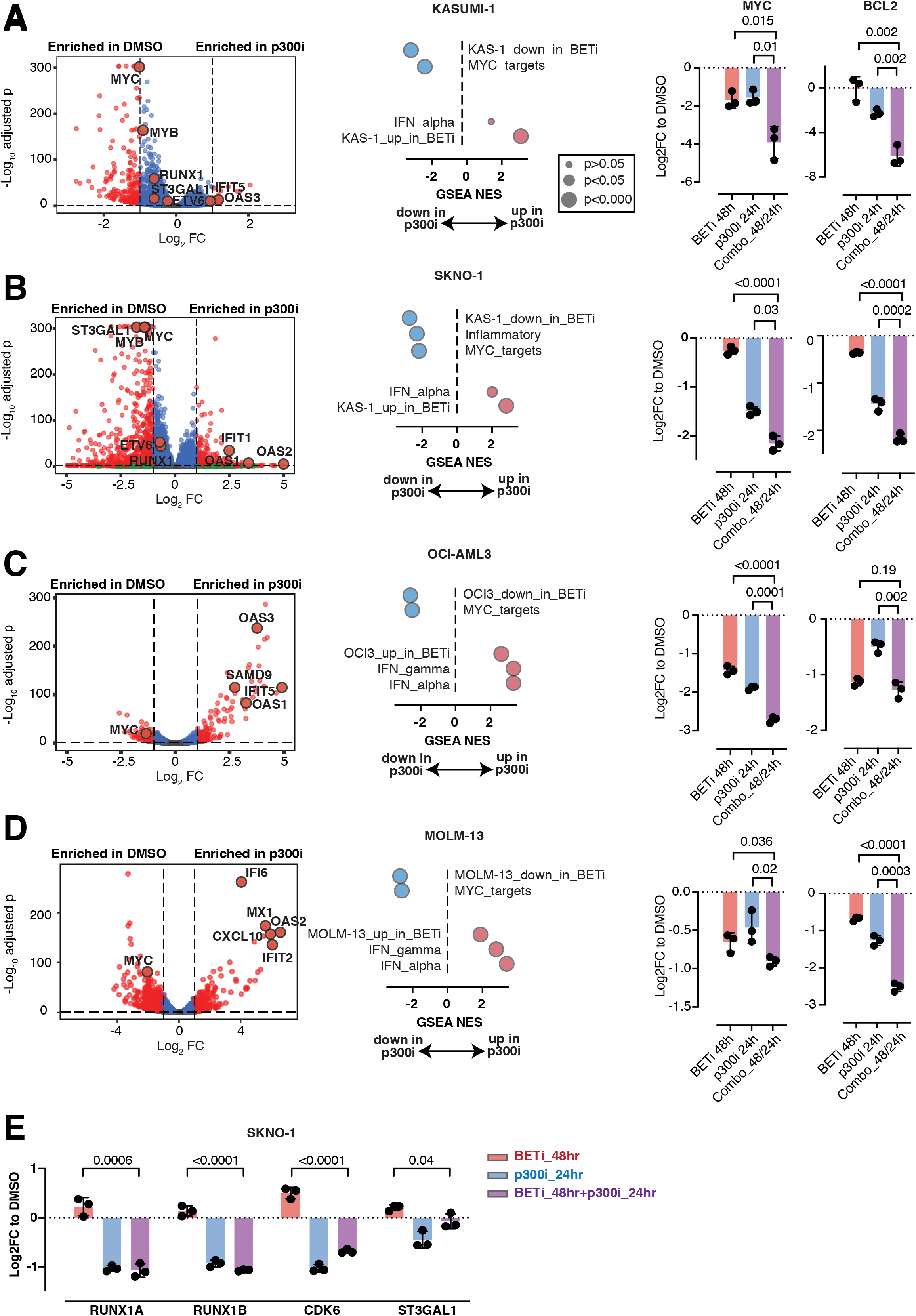
Sequential BETi p300i treatment counteracts expression of rescue programs in a cell type-specific manner. A.-D. Left panels - Volcano plots showing RNASeq expression changes in the indicated cell lines after 24h of treatment with DMSO or p300i. Middle panels - Dot plots of GSEA normalized enrichment scores (NES) and FDRq significance of the top-scoring data sets after treatment with either DMSO or p300i for 24h in the indicated cell lines. Right panels - Analysis of qPCR expression of MYC and BCL2 in the indicated AML cell lines after treatment with either DMSO (48h of treatment), BETi (48h), p300i (24h) or sequential BETi and p300i (48h/24h). Shown are log2 Fold Changes normalized to DMSO-treated controls and SD from 3 biological replicates. E. Analysis of qPCR expression of the indicated transcripts/genes in SKNO1 cells after treatment with either DMSO (48h of treatment), BETi (48h), p300i (24h) or sequential BETi and p300i (48h/24h). Shown are log2 Fold Changes normalized to DMSO-treated controls and SD from 3 biological replicates.

We also performed RNASeq in KASUMI-1 and MOLM-13 cells treated with either DMSO or BETi and combined these with results from our previously reported OCI-AML3 microarray data^12^. Aiming to get a non-biased view of co-dependencies of p300i and BETi, we created specific gene sets of differentially up- or downregulated genes for each of KASUMI-1, OCI-AML3 and MOLM-13 cells following BETi (Supplementary Table S3). We integrated these genes into the Hallmark collection from the Molecular Signatures Database^41^ and finally performed GSEA after treatment with p300i. As KASUMI-1 and SKNO1 behaved identically upon BET inhibition, the gene expression datasets after BETi in KASUMI-1 were also utilized for SKNO1. Comparing these datasets demonstrated that all BETi up- or downregulated gene datasets were significantly enriched in their corresponding p300i-treated counterparts, and mostly consisted of decreased MYC transcriptional targets and increased interferon responsive genes (middle panels Figures 4A-D).

These results suggested a central role for MYC and its transcriptional program as a common target of both BETi and p300i. We hypothesized that a redundant role for BET and p300 in the maintenance of critical AML genes might define the acute resistant state and predict for p300i sensitivity in this state. We determined the effects of sequential BETi followed by p300i treatment on the transcription of *MYC* and the MYC-dependent gene *BCL2* across all 4 AML cell lines. Single agent treatment with either BETi or p300i induced strong downregulation of both genes in all cell lines. However, combined sequential treatment generated significantly stronger downregulation of *MYC* in all, and of *BCL2* in all but one (OCI-AML3) cell line (right panels Figures 4A-D). We also utilized a previously established MYC reporter assay in the unrelated colon cancer HCT116 cell line. Again, combined treatment significantly repressed MYC activity compared with single treatment (Figure S4A). Finally, we determined the effects of sequential BETi followed by p300i on the transcription of the rescue genes *RUNX1_A, RUNX1_B, ETV6* and *ST3GAL1* in SKNO-1 and KASUMI-1 cells. Whereas BETi triggered upregulation of these genes, single p300i and combined sequential treatment led to a downregulation of these transcripts (Figure 4E and S4B).

Altogether, these results demonstrate the dependency of genes that can rescue the apototic phenotype following BETi upon p300i. These results also suggest putative clinical utility of sequential BET inhibition followed by p300 inhibition to specifically and fatally impair transcription of rescue programs such as MYC, thus allowing greater inhibition of leukemic cells.

### p300i is invariably therapeutically effective during early stages of resistance to BET inhibition

Relapsed disease usually presents after many months in AML and whether the mechanisms of acute resistance/persistence still exist at later timepoints is unknown. To address this, we experimentally simulated the evolution of acquisition of resistance to BETi in the two AML models that we exhaustively tested so far: SKNO1 cells to represent t(8:21) AML and OCI-AML3 for *NPM1*mut AML. We treated SKNO1 and OCI-AML3 cells with DMSO or increasing dosages of BETi at IC20 (SKNO1 only), IC40, IC50, IC65, IC80 (SKNO1 only) and IC90 concentrations to generate cell lines along a resistance continuum from early to established resistance (Figure 5A). These stages of resistance evolution were defined based on current growth characteristics and behavior when exposed to the next BETi dose stage. For example, OCI-AML3 BETi IC65_resistant cells had similar growth after 90 (IC65_short_resistant) and 180 days (IC65_long_resistant) in culture. However, increasing BETi dosage to IC90 induced rapid cell death in the IC65_short_resistant culture, whereas the IC65_long_resistant cells remained cellularly resistant, thus allowing us to suggest IC65_long_resistant cells as the first stage of established resistance to BETi. In contrast, IC65_resistant SKNO1 cells quickly died when exposed to IC90 dosages even after 180 days in culture, and required an intermediate dosage (IC80) to overcome the transition to IC90-resistance (data not shown).

**Figure 5.**
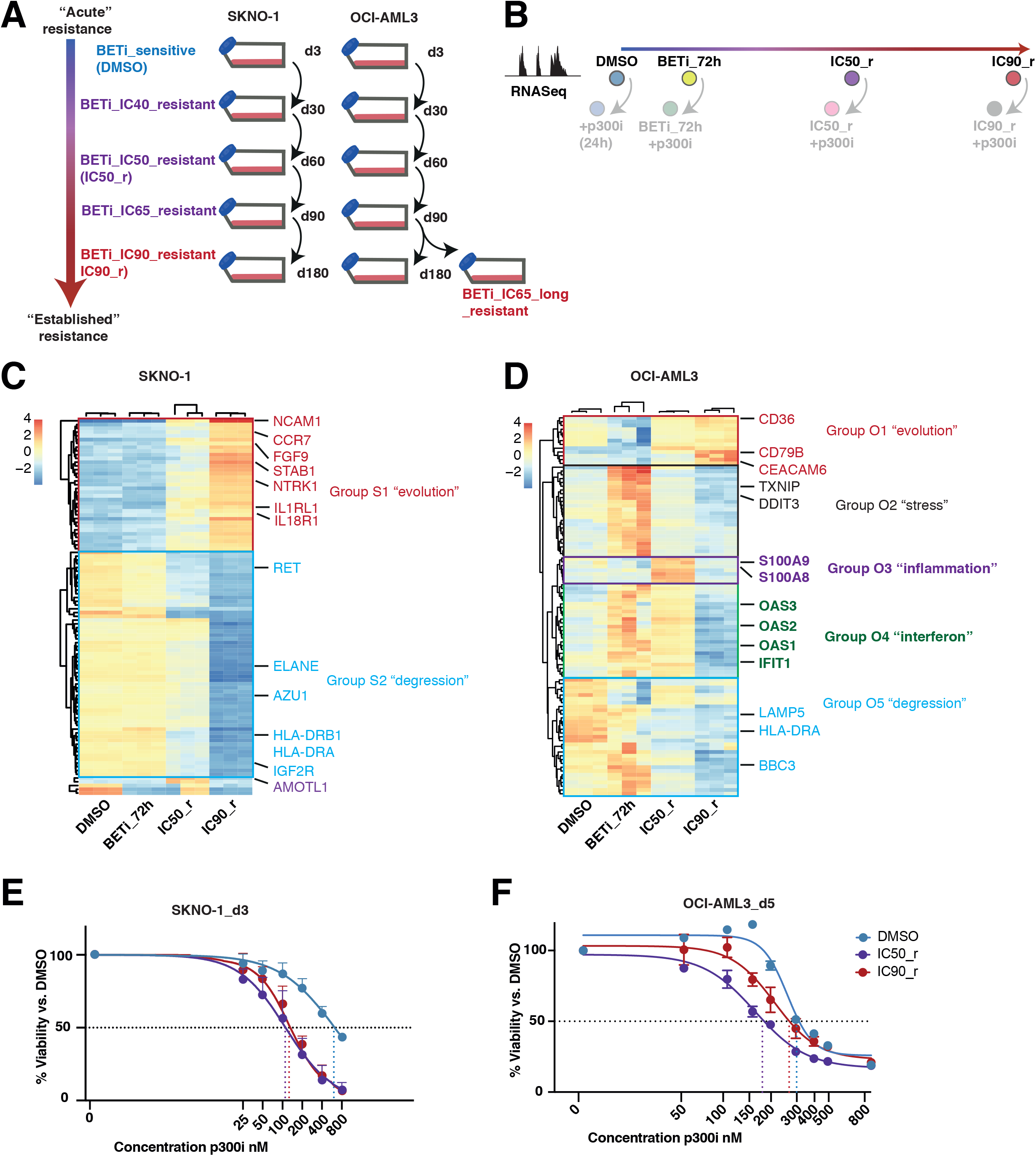
p300i is invariably effective during early stages of resistance to BET inhibition. A. Graphical schema of the experimental approach to induce and store cells at all stages of resistance to BETi in SKNO1 and OCI-AML3 cells. B. Experimental approach to the assessment of transcriptional changes via RNASeq during the longitudinal scale of establishment of resistance to BETi. To avoid putative sequencing-related batch effects, these experiments were directly performed in conjunction with matched p300i-treated (24h) samples (shown with lower opacity). All RNASeq experiments were performed on 3 biological replicates (altogether 24 RNASeq samples per model). C. -D. Variance by non-supervised hierarchical clustering of the 100 most variable genes in DMSO, BETi_72h, IC50_r and IC90_r isogenic SKNO1 (5C) and OCI-AML3 (5D) cells. E.-F. Assessment of cell proliferation of the indicated isogenic SKNO1 (5E) and OCI-AML3 (5F) cell lines after 72h (for SKNO1) and 120h (for OCI-AML3) of p300i treatment. Shown are mean percentages normalized to DMSO-treated controls and SD from 3 biological replicates.

We assessed serial “snap-shots” of transcription during the evolution of resistance to BETi by performing RNASeq at 4 longitudinal time-points: in DMSO-treated, short-term BETi-treated (BETi_72h), incipiently resistant cells (IC50_r) and cells with full-blown resistance to BETi (IC90_r) (Figure 5B). We then performed non-supervised clustering analysis of the most variable genes during the longitudinal process of resistance. The transcriptional patterns of resistance appeared different in the two models. In SKNO1, resistance was binary, with ostensibly only two transcriptional patterns. Group S1 we named “evolution”, because it consisted of genes that continuously increased their expression during resistance (Figure 5C). This group was enriched for genes encoding signaling ligands and receptors such as *NCAM1, CCR7, FGF9, IL1RL1* and others (Figures 5C, 6B). Group S2 we named “degression” and consisted of genes whose expression gradually reduced from levels in untreated cells during the evolution of resistance (Figure 5C). By contrast, in OCI-AML3 cells, the transcriptional evolution of resistance was more complex and certainly non-linear. Here, we identified 5 transcriptional groups that separated according to their dynamics during resistance (Figure 5D). Besides a small “evolution” signature (Group O1) and a larger “degression” group (Group O5), some genes only increased transiently at lower IC doses of BETi (termed by us as “stress” genes - Group O2) (Figure 5D). Of importance, a small number of “inflammation” - associated genes (Group O3) were transiently expressed only in incipiently resistant IC50_r cells and consisted of a network of potential resistance modulators such as S100A9/S100A8^42–44^ (Figures 5D, 6D). Finally, we identified an “interferon” gene group (O4), which demonstrated increased expression immediately after BETi treatment and in IC50_r but were subsequently depleted in IC90_r cells (Figures 5D, 6G).

**Figure 6.**
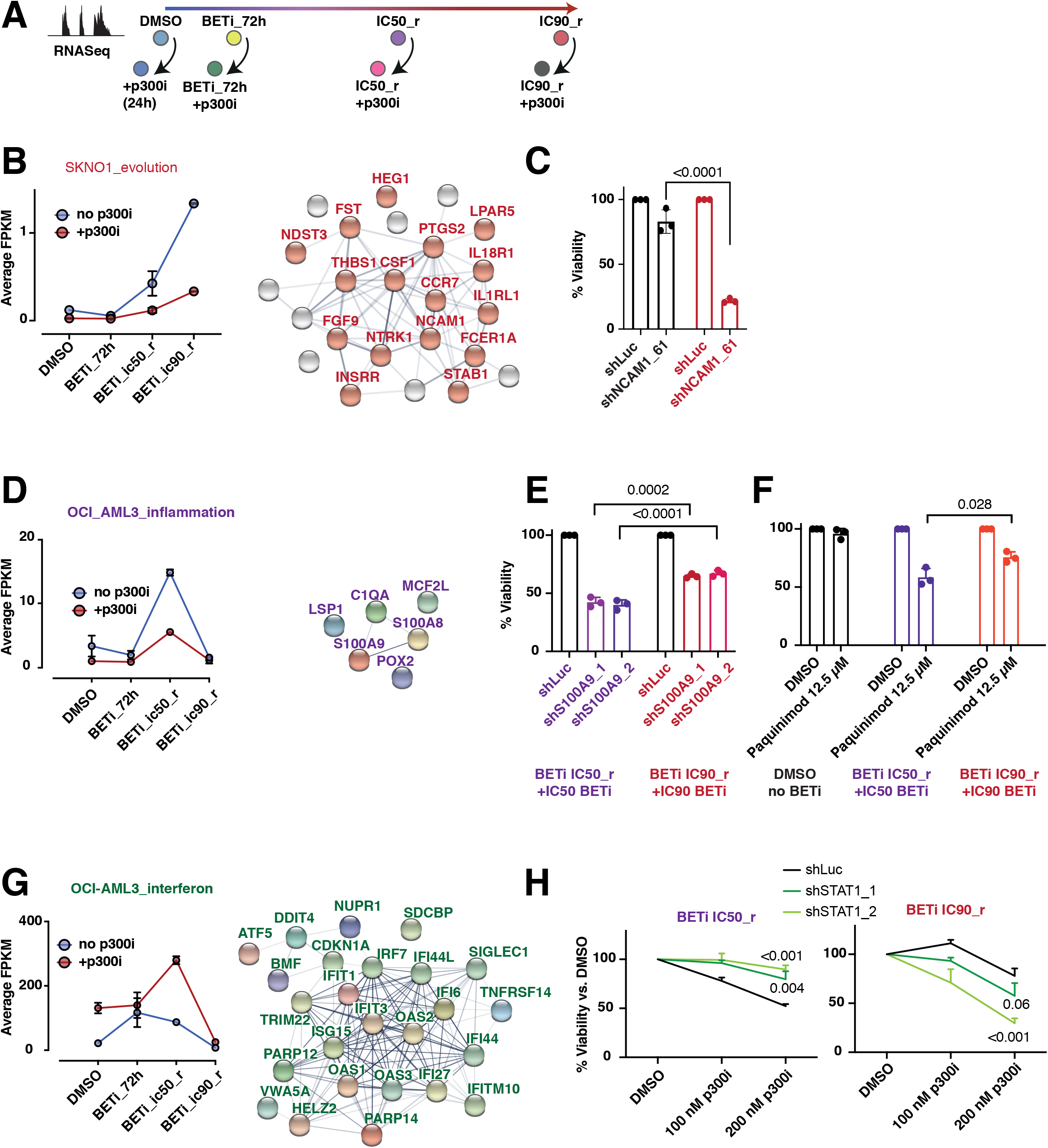
p300 controls multiple routes of resistance to BET inhibition. A. Continued from Figure 5B - Experimental approach to the assessment of transcriptional changes during the longitudinal scale of establishment of resistance to BETi and sensitivity towards p300i. B. Left panel - Longitudinal analysis of expression of a subgroup of SKNO1 evolution genes that were enriched for ligands and signaling receptors, during all stages of resistance to BETi, with or without addition of p300i for 24h. Shown are average FPKM values from 3 biological replicates and SD. Right panels - Graphical depiction and pre-described interactions of all members of the indicated group. C. Assessment of cellular viability after *NCAM1* knockdown in the indicated SKNO1 isogenic cells for 120h. Shown are percentages normalized to Luc controls and SD from 3 biological replicates. D. Left panel - Longitudinal analysis of expression of all OCI-AML3-related “inflammation” genes, during all stages of resistance to BETi, with or without addition of p300i for 24h. Shown are average FPKM values from 3 biological replicates and SD. Right panels - Graphical depiction and pre-described interactions of all members of the indicated group. E. Assessment of cellular viability after S100A9 knockdown in the indicated isogenic cells for 120h. Shown are percentages normalized to Luc controls and SD from 3 biological replicates. F. Analysis of cellular viability after treatment with either DMSO or Paquinimod in the indicated OCI-AML3 isogenic cells for 72h. Shown are percentages normalized to DMSO-treated controls and SD from 3 biological replicates. G. Left panel - Longitudinal analysis of expression of all OCI-AML3-related “interferon” genes, during all stages of resistance to BETi, with or without addition of p300i for 24h. Shown are average FPKM values from 3 biological replicates and SD. The massive boost upon p300i at IC50_resistance should be noted. Right panels - Graphical depiction and pre-described interactions of all members of the indicated group. H. Analysis of cellular viability after STAT1 knockdown and DMSO- or p300i-treatment for 120h in the indicated isogenic cell lines. Shown are percentages normalized to DMSO and SD from 3 biological replicates.

We next treated cells from all stages of BETi-resistance with p300i and measured proliferation. In SKNO1, BETi-resistance severely sensitized cells to p300i throughout all stages (compared with DMSO controls) (Figures 5E, S5A-B). Sensitivity to p300i was highest starting with IC50_r and remained at comparable values during later stages of established resistance, including in IC90_r cells (Figures 5E, S5A-B). In OCI-AML3 cells, although p300i abrogated proliferation at lower IC50 values during all steps of resistance to BETi compared with DMSO controls (Figures 5F, S5C-D), compared with earlier stages of resistance, cells with established resistance to BETi (IC90_r cells) required significantly more p300i to achieve the same inhibitory effects (Figures 5F, S5C-D).

These results suggest cell type/genotype-dependent transcriptional mechanisms of resistance adaptation, yet also reveal an early but relatively wide window of opportunity to improve treatment responses to BETi with the addition of p300i. As sensitivity to p300i was always high during earlier stages of resistance, the data would suggest to introduce the combination prior to the development of established resistance.

### p300 controls multiple routes of resistance to BET inhibition

We next assessed gene expression alterations following p300 inhibition at key stages of adaptation to BETi (DMSO, 72h BETi, IC50_r and IC90_r) (Figure 6A). This experiment aimed to demonstrate whether expression of the most differentially expressed gene groups altered during the evolution of resistance were dependent on p300.

In SKNO1, p300i severely repressed the expression of a group of “evolution” genes, which were enriched for signaling receptors and ligands, in both IC50_r and IC90_r cells (Figure 6B, S6A). The top-scoring “evolution” gene, and likewise one that was significantly re-repressed by p300 inhibition, was *NCAM1* (Figure 5C, 6B). We have previously demonstrated that the aberrant expression of NCAM1 (also known as CD56) leads to resistance in diverse types of AML^45^. Furthermore, NCAM1 is usually expressed and also associates with higher relapse rates in t(8:21) AML^46,47^. To show a direct role for NCAM1 in the development of resistance to BETi in SKNO1 cells, we performed RNAi-mediated knockdown in DMSO-vs IC90_r cells. As shown in Figure 6C, *NCAM1* knockdown severely abrogated cell growth of IC90_r cells, while barely repressing proliferation of DMSO cells.

In OCI-AML3, p300i comparatively repressed the expression levels of “inflammation” genes in IC50_r cells, of “evolution” genes in IC90_r cells, of “stress” genes in short-term BETi-treated cells and those of “degression” genes in general (Figure 6D, S6B). Conversely, p300i massively boosted “interferon” genes in IC50_r cells only (Figure 6G). Given that IC50_r cells were most sensitive to p300i and to further approach the differences between incipient and established stages of resistance in OCI-AML3 cells, we tested if resistance mechanisms at least partly related to upregulation of S100A9, as suggested by the expression of the “inflammation” group (Figure 6D). We either treated cells with Paquinimod, a compound that prevents S100A9 binding to TLR-4 and therefore impairs S100A9 activity, or created cell lines with inducible RNAi-mediated *S100A9* knockdown. In both settings, although depletion of S100A9 function suppressed both cell types, this inhibition was significantly higher in IC50_r than IC90_r cells (Figures 6E-F, S6D).

We also performed inducible RNAi-mediated knockdown of the IFN-effector *STAT1* in IC50_r and IC90_r cells and treated them with p300i. Remarkably, p300 inhibition in *STAT1*-depleted IC50_r cells failed to repress cellular proliferation when compared with DMSO-treated controls. Conversely, p300i treatment of *STAT1*-depleted IC90_r cells significantly suppressed their proliferation (Figure 6H, S6E).

Altogether, even in reductionist systems these data demonstrate the temporal and genotypic complexity of p300-induced adaptation routes to BET inhibition. p300-induced resistance to BETi appears to evolves continuously, and utilizes a number of separate and often transient emergency programs to maintain leukemia fitness along this continuum to established resistance.

## Discussion

The majority of AML patients die, following a period of transient response, from disease relapse^2,48–50^. This suggests an understanding of resistance mechanisms as one major unanswered challenge in leukemia research. Moreover, as resistance mechanisms are likely pleotropic and specific to individual drugs, the importance of this question is amplified by the growing number of promising novel agents available^51,52^. Although many studies have elucidated genetic and non-genetic causes of drug resistance, most have focused on frank relapsed disease, which usually occurs months after initial therapy. However, by definition, resistance must initiate under the selective pressure of the induction therapy^53^ and it remains unknown whether acute mechanisms that underpin initial persistence of leukemia cells are the same as those that prevail in bulk disease upon re-challenge at relapse. This distinction may have clinical implications: response and survival rates are far higher for treatment at diagnosis than in relapse and many physicians feel that for some AML cases, there is only “one-shot” at effecting cure. A greater knowledge of acute mechanisms of resistance could allow rational development of combinations to prevent relapse. We have addressed these questions in our study, using the example of the evolution of BET resistance in susceptible model systems.

Our primary objective was to anticipate chronic resistance by dissecting early routes of adaptation that linked BETi to subsequent cellular persistence. We identified that p300 rapidly ameliorates the loss of transcription at diverse critical AML genes, to explain, at least in part, acute cellular persistence. Whilst our experiments focused on p300 to emphasize the translational consequences and to simplify the narrative, we also observed that RUNX1, RUNX1-RUNX1T1 and FLI1, known critical TFs in KASUMI-1 cells^27,29,30,32,34,54^ were enriched at the top rescue sites, in conjunction with p300. These results reconcile with a demonstrated signaling cascade composed of hematopoietic TFs, p300/CREBBP, and BRD4 to support leukemia maintenance, in which TFs and p300/CREBBP recruit/instruct BRD4 to initiate transcription^36^. However, our data also suggest that an increase in p300 activity can maintain sufficient levels of transcription to allow some cells to persist, even in the presence of reduced BRD4 binding.

It is known that, at some loci, p300 activity is redundant with CREBBP, and their lysine acetyltransferase (KAT) domains share an almost identical homology^38,55,56^. Therefore, it is possible that CREBBP may perform a similar function in resistance to BETi. However this distinction is somewhat academic as p300i inhibit the enzymatic KAT function of both p300 and CREBBP and our proposed therapeutic intervention would also target CREBBP-mediated resistance^38^.

In our model of acquisition of resistance to BETi, synthetic dependency with p300 inhibition differed between models. SKNO1 cells remained at the maximum dependency level between IC50-resistant and IC90-resistant cells. In OCI-AML3, sensitivity towards p300i was highest during earlier stages of resistance and gradually decreased along the continuum to the establishment of full-blown resistance. To determine the mechanism(s) underpinning the development of this dynamic resistance, and its sensitivity to p300 inhibition, we performed longitudinal transcriptional profiling at different stages of resistance. We identified and verified the adhesion molecule NCAM1 as a p300-dependent factor important for resistance in SKNO1 cells. In OCI_AML3, we showed the importance of inflammation-associated S100A factors for early resistance. In addition, we demonstrated upregulation of an IFN-dependent pathway upon p300i that associated with sensitivity of these cells. Taken together, these findings reveal stunning differences between the transcriptional mechanisms and dynamics underpinning resistance between AML subtypes. They reconfirm the resistance-inducing role of NCAM1, an NK cell marker with proven importance in t(8:21) AML and also implicate the inflammatory response and inflammatory signals as important for early resistance in *NPM1mut* cells, adding a new layer of complexity to the current knowledge about transcriptional plasticity-induced resistance to BETi^13,18,22,45,57^. However, while very intriguing, questions remain. Are the inflammatory and interferon signals cell-intrinsic or do they involve autocrine/paracrine communication that might be effective even in our suspension-based systems? Our reductionist systems did not include microenvironment components and it will be important to include these in more sophisticated models in the future. In addition, it remains to be demonstrated why some transcriptional programs are important only during incipient resistance; are they too energy-consuming or do they require residual BRD4 activity that is extinguished at later time points?

Altogether, we demonstrate that acute resistance to BETi occurs through a mechanistic feedback to p300. Under the same selective therapeutic pressures, this acute resistance can then evolve into fully established resistance where p300 continues to regulate temporal- and genotype-specific programs to maintain leukemic fitness. However, our work also establishes the framework for sequential treatment with BETi followed by p300i, that results in an increased early synthetic lethality, thus identifying a rational combination designed to prevent relapse.

## Author contributions

V.S., G.G., H.O., B.J.P.H. and D.S. conceived and designed the experiments. V.S., G.G., H.O., M.M., B.S., H.Y., S.J.H., S.A-S., P.S.H., D.L., F.B. and D.S. performed experiments and formal analysis. D.S. and E.M. performed the bioinformatics analysis. M.W.M.K., B.G., M.T., T.K., P.G., and R.K.P. provided critical resources. D.S. and B.J.P.H. wrote the manuscript draft. All authors reviewed and edited the final manuscript.

## Acknowledgements

This work was supported by a grant of the Else-Kröner-Fresenius-Foundation (2020_EKEA.35) to D. Sasca and grants of Cancer Research UK (CRUK) - C18680/A25508, Kay Kendall Leukaemia Fund (KKLF) - KKL1243, the European Research Council (647685), MRC (MR- R009708-1), Blood Cancer UK, and the Wellcome Trust (205254/Z/16/Z) to B.J.P. Huntly. P. Gallipoli is funded by a CRUK Advanced Clinician Scientist Fellowship (C57799/A27964) and was previously funded by a WT fellowship (109967/Z/15/Z) and ASH global award. Work in the Huntly Lab is supported by the NIHR Cambridge Biomedical Research Centre (BRC-1215-20014), and was funded in part, by the Wellcome Trust who supported the Wellcome - MRC Cambridge Stem Cell Institute (203151/Z/16/Z) and Cambridge Institute for Medical Research (100140/Z/12/Z).

