## Supplemental_Figures_and_Methods for "Acute resistance to BET inhibitors remodels compensatory transcriptional programs via p300 co-activation"

**Supplemental Materials and methods**

**Cell Culture Conditions**

OCI-AML3 (RRID:CVCL_1844), MOLM-13 (RRID:CVCL_2119), and KG-1 (RRID:CVCL_0374) cell lines were cultured in RPMI 1640 (Gibco) supplemented with 10% fetal bovine serum (FBS) (Sigma-Aldrich) and 1% L-glutamine (Gibco). KASUMI-1 (RRID:CVCL_0589) and SKM-1 (RRID:CVCL_0098) were cultured in RPMI 1640 supplemented with 20% FBS and 1% L-glutamine. SKNO-1 (RRID:CVCL_2196) were cultured in RPMI 1640 (Gibco) supplemented with 10% fetal bovine serum (FBS) (Sigma-Aldrich) and 10% conditioned medium of cell line 5637. Cells were expanded for 1 to 3 weeks in liquid culture prior to any experiment and routinely checked for mycoplasma contamination using the Venor GeM Mycoplasma Detection kit. Cell line identity and purity were regularly verified (Multiplexion).

**Primary AML growth culture, drug and proliferation assays**

Human AML mononuclear cells were obtained via Ficoll-PAque PLUS gradient (GE Healthcare) from bone marrow or peripheral blood of patients. Informed consent was obtained in accordance with the Declaration of Helsinki. The study was conducted under authority granting ethics approval (UK Health Departments' Research Ethics Service reference number REC 07-MRE05-44 and Ethik-Kommission der Landesärztekammer Rheinland-Pfalz reference number 837.270.05). Primary AML sample cell culture was performed as previously published [1] with the following remarks. To avoid cell clumps, free DNA was initially enzymatically digested using DNAseI (100 µg/ml) (Thermo) in IMDM (Iscove's modified Dulbecco's medium, Gibco) supplemented with 20% FBS. Afterwards, cells were cultured in IMDM medium supplemented with 15% BIT (bovine serum albumin, insulin, transferrin; Stem Cell technologies Catalog no. 09500), 1 µM ß-Mercaptoethanol, 1 % Penicillin/Streptomycin and 1 µM SR1 (StemRegenin 1, Selleckchem Catalog no. S2858) along with essential cytokines (100 ng/ml human SCF; 50 ng/ml human FLT3-L; 20 ng/ml human IL-3; 20 ng/ml human G-SCF) (all Peprotech, Catalog no. 300-07, 300-19, 200-03 and 300-23, respectively). For treatment assays, cells were cultured in 48-well plates with a final concentration of 5x10^4 cells per well in 500 µL medium for 5-7 days and treated with 2 different concentrations of BETi and p300i and combinations of these. Cellular viability was measured at day 3 and 5 after treatment start using CellTiterGlo 2.0.

**Clonogenic assays in methylcellulose**

Human AML cell lines, human primary AML cells and murine AML cells were plated in duplicate in methylcellulose medium in the presence of DMSO, BETi or p300i. MethoCult H4435 (STEMCELL Technologies) was used for human AML, and MethoCult GF M3434 (STEMCELL Technologies) was used for murine AML. For AML cell lines and for murine Aml1-Eto9a cells, 30,000 cells per plate were used. For primary AML samples, 80,000 cells per plate were used. Replating was performed at 7 days.

**Flow cytometry**

Flow cytometry experiments were performed on either a BD Fortessa or a BD Canto II.

**Chromatin Immunoprecipitation (ChIP), library preparation, sequencing**

For the preparation of ChIP DNA and ChIPSeq libraries please refer to our previous work[2]. Libraries were sequenced on an Illumina HiSeq 2500 for 50 base pairs in single read mode (for KASUMI-1 data) or on an Ilumina NovaSeq 6000 instrument in 150 bp paired-end mode (for all SKNO1 and OCI-AML3 data).

**ATAC-Seq library preparation and sequencing**

For ATAC-Seq libraries, 100,000 cells were pelleted and washed with 500 µl ice-cold PBS, then lysed with 400 µL ice-cold sucrose buffer (10 mM Tris-Cl pH 7.5, 3 mM $CaCl_2$, 2 mM $MgCl_2$ and 0.32 M sucrose + freshly added 0.5% Triton-X 100 after 12 minutes of initial incubation) and re-collected via centrifugation for 10 minutes at 500 G and 4°C. The resulting nuclear pellet was tagged- and fragmented using 20 µL of nuclease-free water and 5 µL Tagment DNA Enzyme 1 (Illumina Catalog no. FC-121-1030) for 30 minutes at 37 °C. The tagmentation reaction was stopped by addition of 500 µL buffer PB. Afterwards, transposed DNA was purified with the MinElute PCR Purification kit (Qiagen) and amplified using the following PCR Cycler conditions: step 1 - 72°C 5 minutes, 98°C 30 seconds; step 2 - 11-12 cycles: 98°C 10 seconds, 63°C 30 seconds, and 72°C 1 minute; step 3 - 72°C 5 minutes. Finally, PCR products were re-purified with the Qiagen MinElute PCR Purification kit. DNA libraries were quantified with the KAPA Library Quantification kit (Roche Catalog no. KR0405). Library quality was checked on a Bioanalyzer 2100 system with High Sensitivity DNA chips (Agilent Technologies, Catalog no. 5067-4626). Libraries were sequenced on an Illumina HiSeq 2500 for 50 base pairs in single read mode. Two biological replicates were performed independently for each cellular condition.

**Vectors, virus production, transfection and transduction**

The MigR1-AE9a (kindly provided by Dong-Er Zhang, RRID:Addgene_12433)[3] retroviral vector and the psiEco packaging plasmid were transfected in a 1:1 ratio into 293T (RRID:CVCL_0063) cells using TransIT-LT1 transfection reagent (Mirus Catalog no. MIR 2306) according to the manufacturer's protocol. Supernatant was harvested 48 and 60h after transfection. For transduction, one million cKIThighTer-119negative fetal liver cells were mixed in a 1:1 ratio with retroviral supernatant supplemented with mIL-3, IL-6, and murine stem cell factor cytokines (final concentrations of 10 ng/ml, 10 ng/ml, and 100 ng/ml respectively; all Peprotech Catalog no. 213-13, 216-16 and 250-03, respectively) and Polybrene (Sigma-Aldrich, Catalog no. TR 1003) to a final concentration of 8 ng/µl. Spinoculation was performed serially for 2 times.

The shRNAs targeting *STAT1, NCAM1* and *S100A9* were chosen from the TRCN Broad Institute Database and cloned into Tet-pLKO.1-puro vectors (kindly provided by Dimitri Wiederschain, Addgene_21915)[4]. Following shRNA target sequences were used:

*shSTAT1_1*: 5'-GAACAGAAATACACCTACGAA-3'

*shSTAT1_2*: 5'-CGACAGTATGATGAACACAGT-3'

*shNCAM1_61*: 5'-GCCAAGCTCCAATTACAGCAA-3'

*shS100A9_1*: 5'-CGCAACATAGAGACCATCATC-3'

*shS100A9_2*: 5'-CATCAACACCTTCCACCAATA-3'

*shLuc_1*: 5'-ATGTTTACTACACTCGGATAT-3'

Lentiviral particles were produced by co-transfection with psPAX2 (Addgene_12260) and pMD2.G (Addgene_12259) in 293T cells using the transfection reagent TransIT LT-1. Spinoculation of lentivirus was performed one time in the presence of 5 µg/mL Polybrene. Cells were selected with 1.5 µg/mL Puromycin (Sigma-Aldrich).

**Animals and bone marrow transplantation**

CKIT-high/Ter-119-negative fetal liver cells were collected from C57BL/6J (B6) (MGI:3028467) mice at E12.5-E14.5. as previously published[3]. Transduced cells were either transplanted into 8-to-10-week old lethally irradiated B6 mice via tail-vein injection or plated into methylcellulose and serially replated for at least 4 rounds. For in-vivo treatments with BETi or vehicle control, secondary transplantations into sub-lethally irradiated age-matched B6 mice were undertaken. Treatment was performed via chow diet supplemented with BETi (180 mg/kg) (GlaxoSmithKline) or a control compound starting at day +13 post-transplantation. Mice were sacrificed at first signs of disease. All animal experiments complied with local and UK national regulations and were performed under an existing UK Home Office project licence (PA46C00DB).

**Immunohistochemistry**

All murine bone marrow, spleen and liver tissues were fixed, embedded and prepared as described previously[5]. In short, tissues were fixed in a 10% formalin solution (CellPath Ltd.). Bones were decalcified in a 0.38 M EDTA (pH7) solution. Tissue sections (4 µm) were stained with Hematoxylin and Eosin (Thermo) according to the manufacturer's protocol.

**Oligonucleotides**

Following qPCR primers were used:

*BCL-2*_fwd: 5'-ATGTGTGTGGAGAGCGTCAA-3'

*BCL-2*_rev: 5'-TTCAGAGACAGCCAGGAGAAA-3'

*MYC*_fwd: 5'-ATGAGGAGACACCGCCCA-3'

*MYC*_rev: 5'-GGAGCCTGCCTCTTTTCCAC-3'

*STAT1*_fwd: 5'-AACAACCACACGGGGTAGG-3'

*STAT1*_rev: 5'-TCTCCCCAAATGTCCCTATTT-3'

*S100A9*_fwd: 5'-CGGCTTTGACAGAGTGCAAG-3'

*S100A9*_rev: 5'-GCCCCAGCTTCACAGAGTAT-3'

*CDK6*_fwd: 5'-TGCAGGGAAAGAAAAGTGCAATG-3'

*CDK6*_rev: 5'-TCCTCGAAGCGAAGTCCTCA-3'

*ST3GAL1*_fwd: 5'-GACTTGGAGTGGGTGGTGAG-3'

*ST3GAL1*_rev: 5'-ACAAGTCCACCTCATCGCAG-3'

*ST3GAL4*_fwd: 5'-GGGAGATGCCATCAACAAGT-3'

*ST3GAL4*_rev: 5'-GGAAGTCCATTGCCTTGAAA-3'

*RUNX1*_fwd: 5'-ACTCGGCTGAGCTGAGAAATG-3'

*RUNX1*_rev: 5'-GACTTGCGGTGGGTTTGTG-3'

*RUNX1*_A_fwd: 5'-GAACCACTCCACTGCCTTTAAC-3'

*RUNX1*__A_rev: 5'-ATTCTGAGGGCTGTCATCTTTC-3'

*RUNX1*_B_fwd: 5'-TGCATGATAAAAGTGGCCTTGT-3'

*RUNX1*_B_rev: 5'-CGAAGAGTAAAACGATCAGCAAAC-3'

*RUNX1*_C_fwd: 5'-TGGTTTTCGCTCCGAAGGT-3'

*RUNX1*_C_rev: 5'-CATGAAGCACTGTGGGTACGA-3'

**Critical antibodies**

Following antibodies were used for ChIP:

Rabbit Anti-H3K27ac Polyclonal Antibody, Abcam Cat# ab4729, RRID:AB_2118291

Rabbit anti-AML1-ETO Polyclonal Antibody, Diagenode Cat#C15310197, RRID:AB_2891230

Rabbit anti-RUNX1 Polyclonal Antibody, Abcam Cat#ab23980, RRID:AB_2184205

Rabbit anti-PU.1 Polyclonal Antibody (T-21), Santa Cruz Biotechnology Cat#sc-352, RRID:AB_632289

Rabbit anti-p300 Polyclonal Antibody (C-20), Santa Cruz Biotechnology Cat# SC-585, RRID:AB_2231120

Mouse anti-p300 Monoclonal antibody, Abcam Cat# ab14984, RRID:AB_301550

Goat anti-LMO2 Polyclonal Antibody, R and D Systems Cat# AF2726, RRID:AB_2249968

Rabbit anti-FLI1 Polyclonal Antibody, Abcam Cat# ab15289, RRID:AB_301825

Rabbit anti-ERG Polyclonal Antibody (C-17), Santa Cruz Biotechnology Cat# sc-354, RRID:AB_2098432

Rabbit anti-CDK9 Polyclonal Antibody, Bethyl Cat# A303-493A, RRID:AB_10949230

Rabbit anti-BRD4 Polyclonal Antibody, Bethyl Cat# A301-985A100, RRID:AB_2620184

Rabbit anti-BRD4 Polyclonal antibody, Diagenode Cat# C15410337

**Nuclear isolation for nuclear RNASeq**

Enrichment and extraction of nuclear RNA was performed as published previously[6].

**RNA-isolation, quantitative Real Time Polymerase Chain Reaction (qRT-PCR) and sequencing**

RNA was isolated with the RNeasy Plus Mini® Kit (Qiagen) according to the manufacturer's instructions. Quantitative real-time PCR was performed on a Agilent Mx3000P Instrument using the Brilliant II SYBR® Green QPCR Master Mix (Agilent) or on a Thermo QuantStudio 5 Analyser (Thermo Fisher Scientific) using the Luna® Universal Probe One-Step RT-qPCR Kit (New England Biolabs).

For RNA-seq experiments, stranded polyA selected libraries were prepared using the TruSeq Stranded mRNA Library Prep Kit® from Illumina according to manufacturer's standard protocol. Libraries were either 125 bp paired-end sequenced on an Illumina HiSeq 2500 or 150 bp paired-end sequenced on an Ilumina NovaSeq 6000 instrument.

**RNA-seq and nucRNA-seq data analysis**

Sequencing reads were quality checked and adapter trimmed using FastQC (RRID:SCR_014583). Reads were aligned to the hg19 human reference genome using STAR (RRID:SCR_004463) two-pass mode (star 2.4.0.1). Transcript-level counts were generated with HTSeq (RRID:SCR_005514) (version 0.6.1p1). Normalization, transformation and differential gene expression analysis were performed using DESeq2 version 1.12.4 (RRID:SCR_015687). Gene enrichment was calculated with GSEAPreranked (RRID:SCR_003199) on all transcripts that were expressed in at least one condition, using "meandiv" as normalization and "weighted" as the scoring scheme. Overrepresentation analysis was performed on defined gene lists with EnrichR (RRID:SCR_001575) using the standard conditions. Volcano plots were performed using the EnhancedVolcano (RRID:SCR_018931) package in R, heatmaps with either the Pheatmap package (RRID:SCR_016418) or with GraphPad Prism (RRID:SCR_002798) Version 8, Venn diagrams with venny (Stefan Jol - *Make a Venn Diagram*. https://www.stefanjol.nl/venny).

**ChIP-seq and ATAC-seq data analysis**

Sequencing reads were quality checked and adapter trimmed using FastQC. Reads were aligned to the hg19 human reference genome using Bowtie2 (RRID:SCR_016368). SAM files were converted into BAM format, sorted, de-duplicated and indexed with Picard (RRID:SCR_006525) and Samtools (RRID:SCR_002105). Reads mapping to repetitive regions or to the mitochondrial genome were discarded. ChIP-seq peaks were called using Macs2 (RRID:SCR_013291), accepting model building and lambda. ATAC-seq peaks were called using Macs2, accepting lambda but not model building. Differential peak enrichment was calculated with the DiffBind (RRID:SCR_012918) package in R.

**Statistics**

Unless otherwise specified, data are presented as mean ± standard deviation (SD). For animal studies, Kaplan-Meier survival analysis was performed and survival was calculated using the log-rank test. Comparisons between two groups were performed using the unpaired Student t test. Comparisons between multiple groups were performed using one-way ANOVA with multiple comparisons. $P<0.05$ was considered significant.

**Supplemental Figure Legends**

**Supplemental Figure S1**

A. Related to Figure 1C - Representative photomicrographs of colony formation of KASUMI-1 and SKNO1 cells after treatment with either DMSO or BETi for 7 days in methylcellulose.

B. Related to Figure 1E - Serial colony formation assays with Aml1-Eto9a-transformed $Kit^{+}Ter$-$119^{-}$ fetal liver cells.

C. Related to Figure 1E - Representative flow cytometry plot from a recipient animal, transplanted with Aml1-Eto9a-transformed $Kit^{+}Ter$-$119^{-}$ fetal liver cells (bone marrow).

D. Related to Figure 1E - Representative histology sections from a recipient animal transplanted with Aml1-Eto9a-transformed $Kit^{+}Ter$-$119^{-}$ fetal liver cells, to demonstrate extensive disease dissemination in the bone marrow, liver, spleen, and kidney.

**Supplemental Figure S2**

A. and B. Representative scatter plots showing Annexin V/7-AAD (A) and cell cycle histograms (B) at 24h after treatment of KASUMI-1 and SKNO1 cells with either DMSO or BETi, to demonstrate that cells were viable and showed no change in the cell cycle appearance when compared with DMSO-treated controls. Shown are representative examples from multiple (n>3) biological replicates that were required during the whole project.

C. Related to Figure 2A-B - Tornado plots to show binding differences between BETi and DMSO conditions for the indicated TFs, regulators and further chromatin parameters in KASUMI-1 cells (24h treatment). Positive enrichment (in red) shows stronger binding upon BETi. Negative enrichment, colored in white, shows loss of binding upon BETi. Shown are representative matched replicates. As they were only required for the initial pattern identification, single replicates were acquired for ChIPSeq profiles and 2 biological replicates for nucRNASeq and ATACSeq.

D. Related to Figure 2E - Average binding curve profiles for RUNX1, RUNX1-RUNX1T1, FLI1 and H3K27ac at the top 5% rescued sites in KASUMI-1 cells.

E. More examples of BRD4 and p300 binding profiles in DMSO and BETi-treated KASUMI-1 and SKNO1 cells, to demonstrate the BETi-triggered increase of p300 binding at the indicated genomic loci.

F. Example of BRD4 and p300 binding profiles in DMSO and BETi-treated OCI-AML3 cells, to demonstrate the BETi-triggered increase of p300 binding at the indicated genomic locus.

**Supplemental Figure S3**

A. Related to Figure 3E - Three-dimensional diffusion plots of synergy of combined treatment with BETi and p300i in the indicated orders and cell lines. Treatment efficiency was measured with CellTiterGlo® at day 5 after commencement of the first treatment. For the sequential treatment modes, the second compound was added 48 hours after treatment commencement. Shown are averages from 3 biological/experimental replicates.

B. Related to Figure 3H - Individual plots showing treatment efficiency of concomitant or sequential treatment with BETi and p300i in 5 primary AML patient samples. Treatment efficacy was measured with CellTiterGlo® at day 4 after commencement of the first treatment. Shown are results from 2 biological replicates for each primary sample. For the sequential treatment mode with BETi first, p300i was added 48 hours after treatment commencement.

**Supplemental Figure S4**

A. Related to Figures 4A-D - Analysis of Luciferase activity, corresponding to the MYC function, in HCT-116-Luc-MYC cells after treatment with either DMSO (48h of treatment), BETi (48h), p300i (24h) or sequential BETi and p300i (48h/24h). Shown are mean percentages normalized to DMSO-treated controls and SD from 3 biological replicates.

B. Related to Figure 4E - Analysis of qPCR expression of the indicated transcripts/genes in KASUMI-1 cells after treatment with either DMSO (48h of treatment), BETi (48h), p300i (24h) or sequential BETi and p300i (48h/24h). Shown are log2 Fold Changes normalized to DMSO-treated controls and SD from 3 biological replicates.

**Supplemental Figure S5**

A. and C. Assessment of cell proliferation of the indicated isogenic SKNO1 (A.) and OCI-AML3 (C.) cell lines after 72h (for SKNO1) and 120h (for OCI-AML3) of p300i treatment. Shown are mean percentages normalized to DMSO-treated controls and SD from 3 biological replicates.

B. and D. IC50 values for p300 inhibition in the indicated cell lines along the stages of resistance to BETi.

**Supplemental Figure S6**

A.-B. Longitudinal analysis of expression of the indicated groups of genes in SKNO1 (A.) and OCI-AML3 (B.) cells, during all stages of resistance to BETi, with or without addition of p300i for 24h. Shown are average FPKM values from 3 biological replicates and SD.

C.-E. Related to Figures 6C, 6E and 6H - Analysis of transcript knockdown efficiency of NCAM1 in SKNO1 (C.), S100A9 (D) and STAT1 (E) in OCI-AML3 cells along the indicated stages of resistance to BETi.

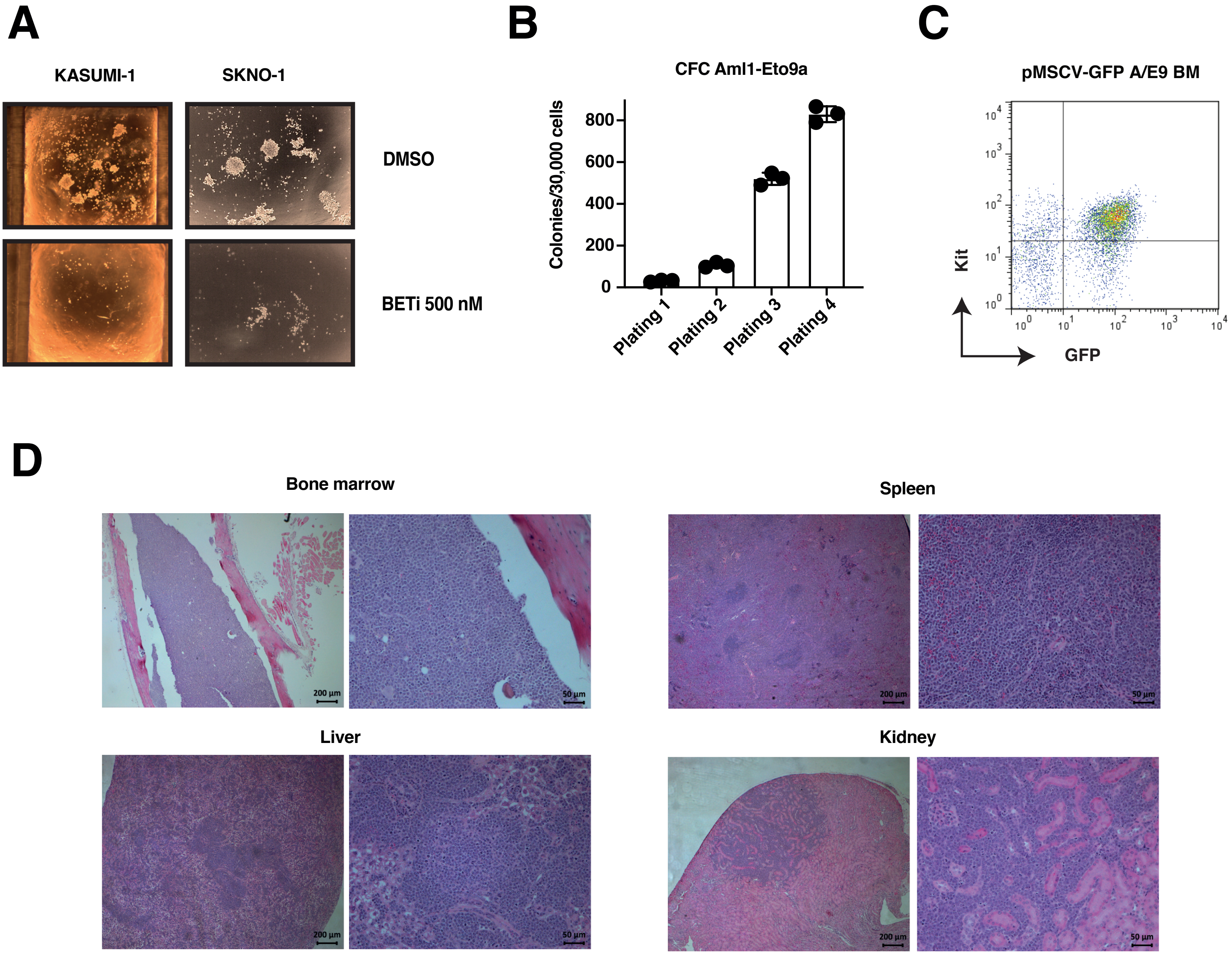


Suplemental Figure S1

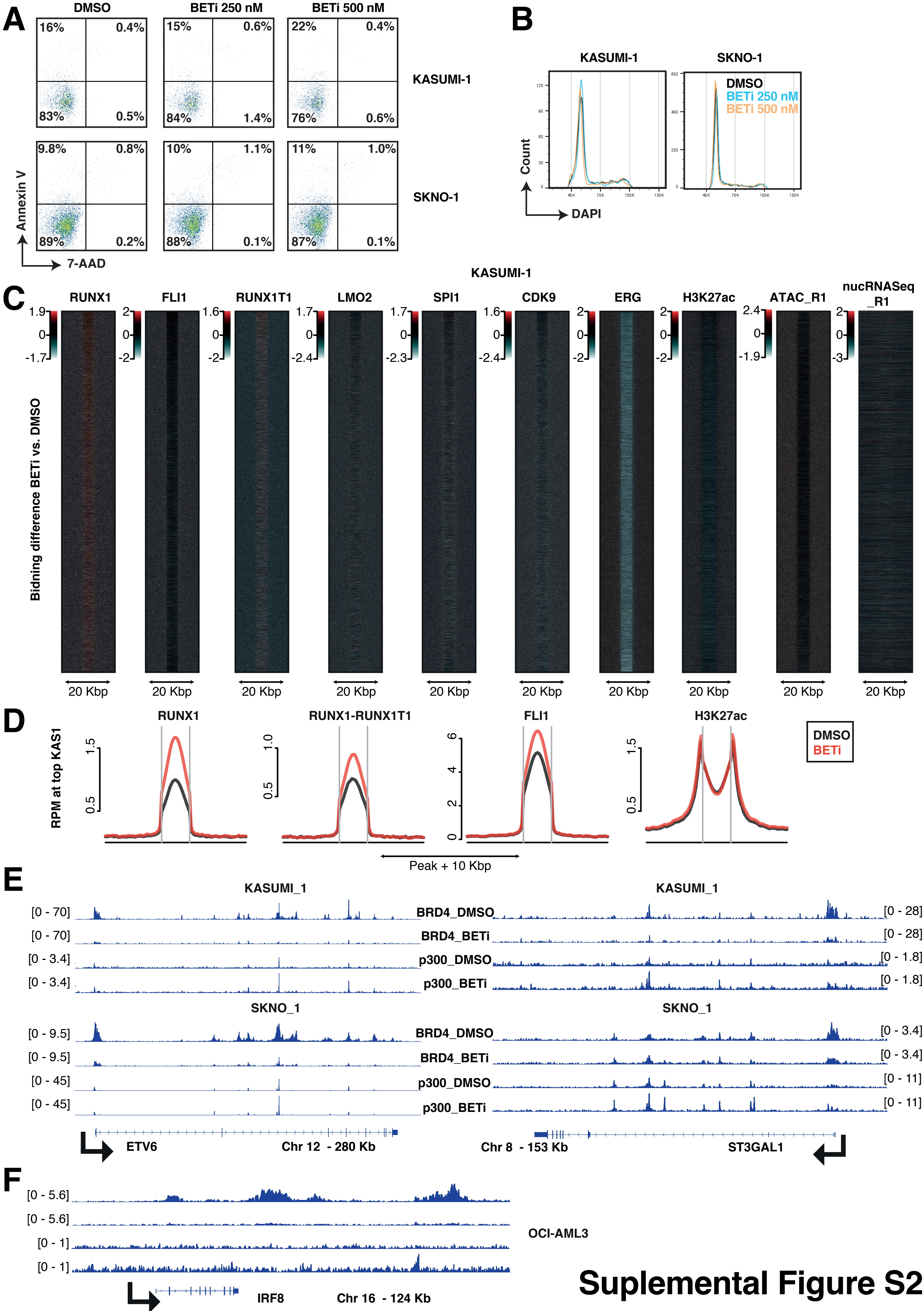


Suplemental Figure S2

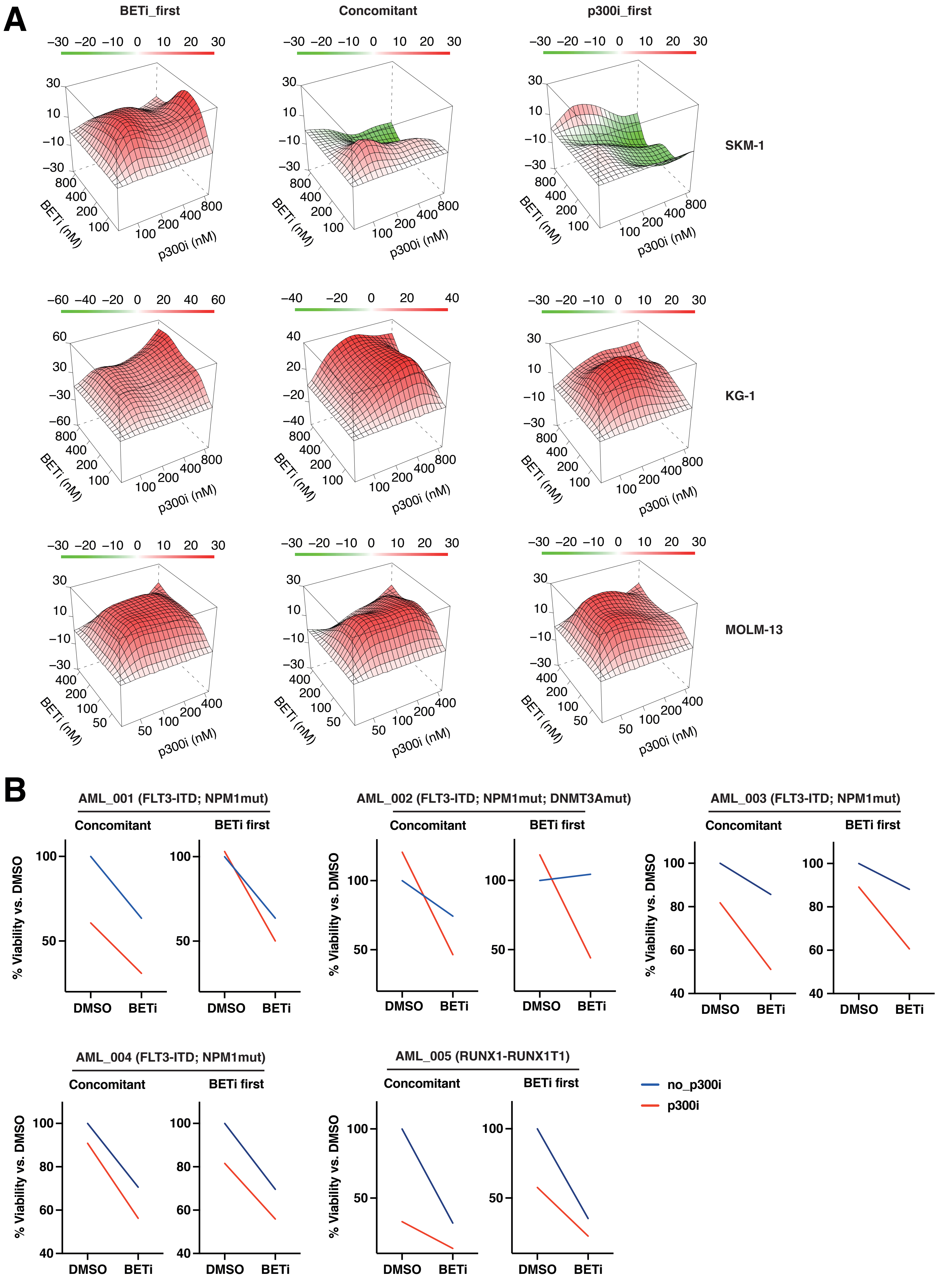


**Suplemental Figure S3**

A

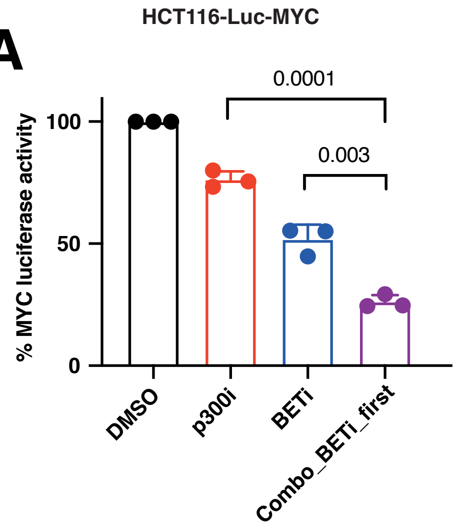


B

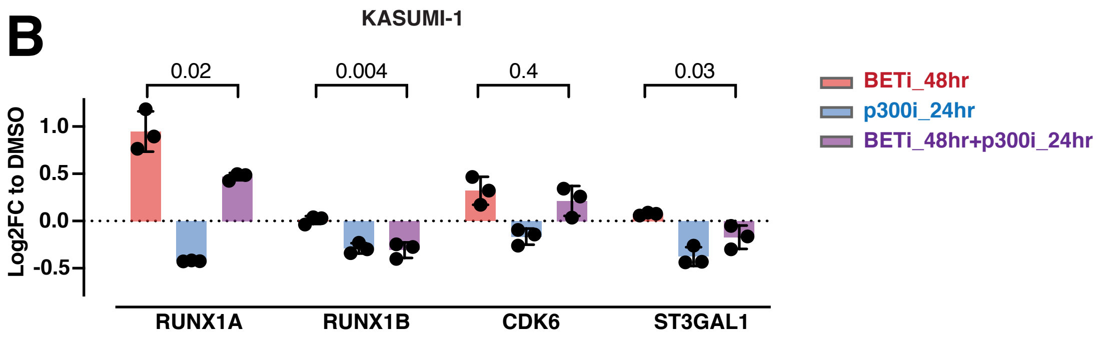


**Supplemental Figure S4**

A

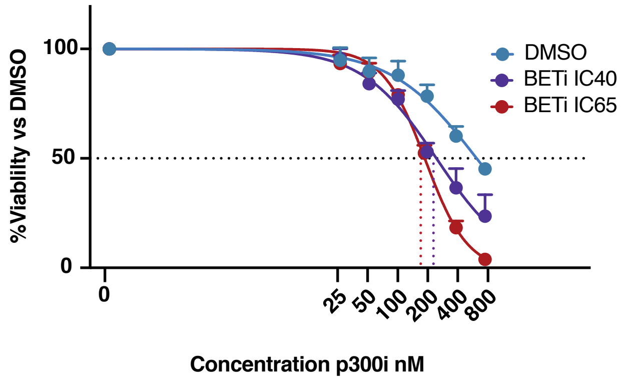


B

p300i nM

| | |
|---|---|
| DMSO | 616 nM |
| IC40_r | 223 nM |
| IC50_r | 110 nM |
| IC65_r | 182 nM |
| IC90_r | 123 nM |

C

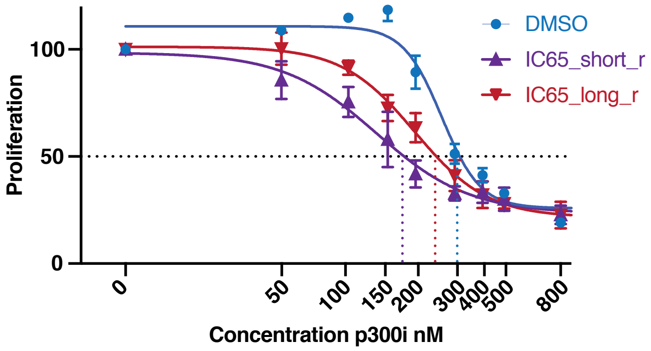


D

p300i nM

| | |
|---|---|
| DMSO | 304 nM |
| IC50_r | 177 nM |
| IC65_short_r | 173 nM |
| IC65_long_r | 247 nM |
| IC90_r | 275 nM |

### Supplemental Figure S5

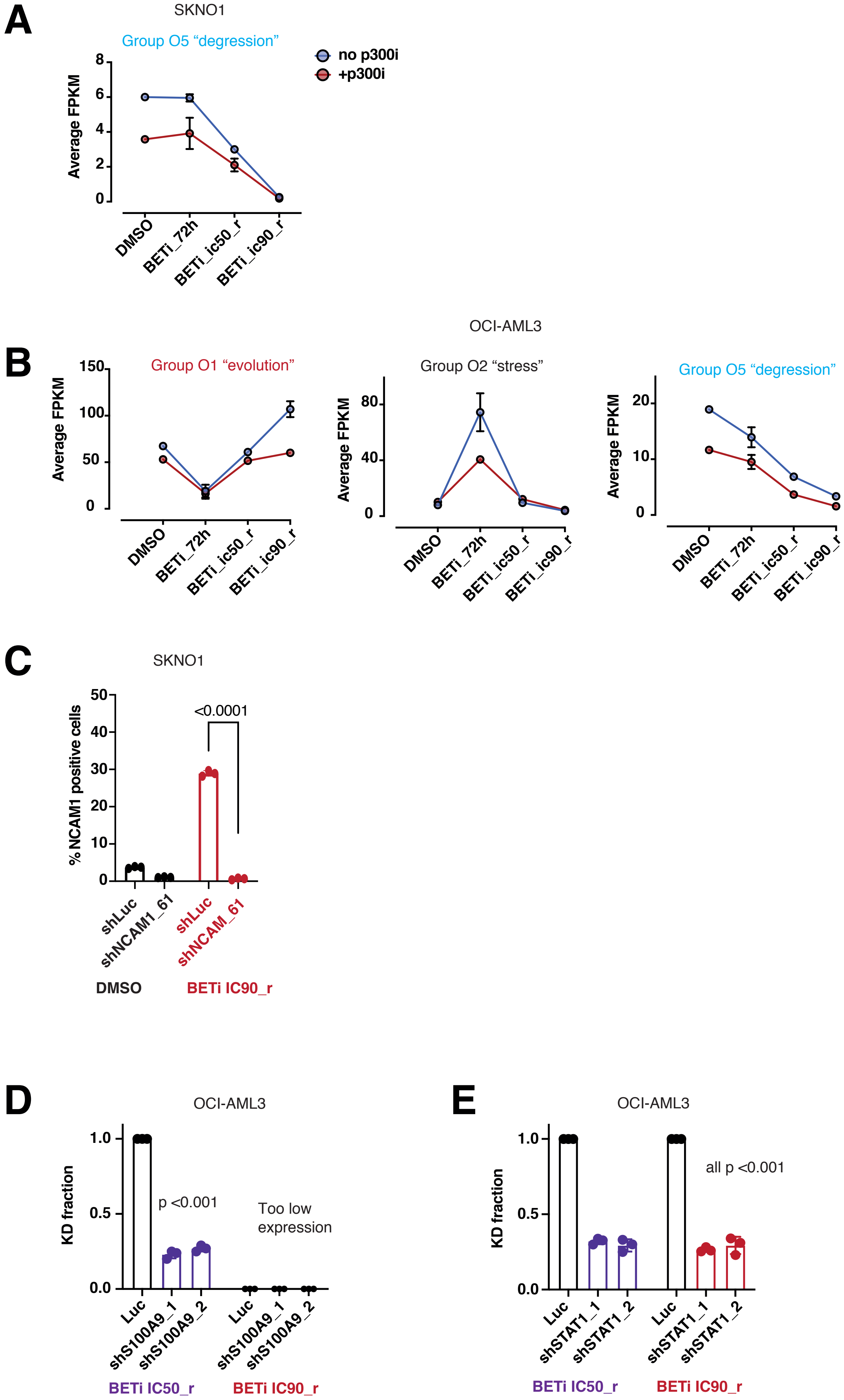


Supplemental Figure S6
